# Unified Probabilistic Analysis of CyTOF: A Deep Generative Approach using CytoOne

**DOI:** 10.64898/2025.12.03.692122

**Authors:** Yuqiu Yang, Kaiwen Wang, Yike Shen, Jon A Weidanz, Guanghua Xiao, Xinlei Wang

**Affiliations:** Quantitative Biomedical Research Center, Department of Health Data Science and Biostatistics, Peter O’Donnell Jr. School of Public Health, University of Texas Southwestern Medical Center, Dallas, TX, USA, 75390; Department of Data Science, Davidson College, Davidson, NC, USA, 28035; Department of Earth and Environmental Sciences, University of Texas at Arlington, Arlington, TX, USA, 76019; Department of Biology, University of Texas at Arlington, Arlington, TX, USA, 76019; Department of Bioinformatics, University of Texas Southwestern Medical Center, Dallas, TX, USA, 75390; Simmons Comprehensive Cancer Center, University of Texas Southwestern Medical Center, Dallas, TX, USA, 75390; Department of Mathematics, University of Texas at Arlington, Arlington, TX, USA, 76019; Division of Data Science, College of Science, University of Texas at Arlington, Arlington, TX, USA, 76019

## Abstract

Extracting meaningful biological signals from Cytometry by time-of-flight (CyTOF) data remains challenging due to heterogeneity, data characteristics, and the presence of various technical artifacts. Current analysis workflows typically rely on task-specific tools assembled into pipelines, which often make inconsistent distributional assumptions and fail to fully leverage the structure of the data. We present CytoOne, a unified probabilistic framework tailored for CyTOF data that integrates batch correction, differential analysis, and visualization within a single model. CytoOne is built upon a Bayesian hierarchical architecture inspired by Nouveau Variational Autoencoders (NVAE) and employs a novel quasi zero-inflated softplus-normal (QZIPN) likelihood to flexibly model the sparse and noisy nature of CyTOF measurements. We demonstrate via qualitative and quantitative evaluations that CytoOne effectively approximates both marginal and joint distributions of CyTOF data, removes batch-specific artifacts, enables fine-grained differential expression analysis, and facilitates interpretable embeddings for exploratory analysis.

## Main

As one of the most commonly adopted proteomic techniques, cytometry by time-of-flight (CyTOF) is capable of simultaneously profiling expressions of up to 120 proteins presented on the surface of tens of thousands or even millions of cells at a single-cell resolution in one single experimental run (1). When coupled with meticulous experimental designs and observed phenotypes, the abundant information such as cell types, cellular differentiation, and differential in expressions contained in CyTOF data provides invaluable insights into the human immune system (2–4).

However, as with most single-cell techniques, reliably extracting said biological signals and thus offering interpretation over the large volume of protein measurements remains a daunting task. First of all, as the exploratory data analysis conducted in (5) has revealed, because of the intricate internal data manipulation automatically carried out during a CyTOF experiment (6), the data distribution of protein measurements does not conform to any basic probability distributions like the Gaussian distribution or the negative binomial distribution that researchers often exploit in the context of Flow Cytometry (7) and scRNA-seq (8, 9). What’s more, due to the existence of the very biological signals that we wish to unveil, the data is of highly heterogeneous nature, rendering the distribution a complex mixture of non-trivial high-dimensional distributions with convoluted dependency relationships (5). To make matters even worse, the measured signals are often contaminated by various artifacts such as batch effects and signal drifts (10, 11).

In this endeavour, a variety of data analysis tools have been devised, aiming either to remove unwanted variation, facilitate biological discovery, or visualize data (10, 12). As extensively reviewed in (5), such tasks generally involve batch effect removal (12, 13), differential discovery (14, 15), visualization (13–17) etc. Powerful as those algorithms are, they are often proposed to solve one specific task, compelling researchers to resort to a pipeline workflow where different methods are heuristically assembled in a sequential manner to generate desired outcomes (16, 17). Intuitive as they might appear, pipelines often contain methods with conflicting or even erroneous distributional assumptions that are not suitable for CyTOF data. For instance, in (16), after performing cell clustering with FlowSOM (18) which was designed to incorporate the complex distributions of CyTOF data, they carried out differential analysis based on a linear regression with Gaussian noise. The discrepancy among assumptions could confound the results and render the reported significance level questionable.

To address these issues, we propose CytoOne, a unified probabilistic framework with a consistent distributional assumption tailored for CyTOF data that can be further applied to an array of essential data analysis tasks such as visualization via dimension reduction, differential discovery, and batch effect removal. CytoOne is constructed as a Bayesian hierarchical model where the likelihood as well as the posterior distributions are specified using deep neural networks in a similar fashion to VAE (19, 20). The optimization is accomplished via stochastic gradient descent (21) which is particularly suitable for large datasets we encounter in CyTOF experiments. We further carry out numerical validations on dimension reduction, differential discovery, and batch effect removal using both simulation and real data. Via benchmarking our performance for each task against corresponding most commonly used methods, we demonstrate while being designed for realizing multiple functionalities, CytoOne still manages to achieve favorable outcomes for all tasks due to its joint optimization nature.

## Results

### The CytoOne model

The development of CytoOne is motivated by the need for a single coherent probabilistic model capable of performing multiple essential tasks on CyTOF data—namely visualization, batch effect correction, and differential expression analysis. Several probabilistic models based on the framework of variational autoencoders (VAEs) (19) have been proposed for carrying out similar tasks in the context of scRNA-seq data (22, 23). However, as (5) has shown, adapting tools intended for scRNA-seq for the analysis of CyTOF data would be non-trivial and could result in suboptimal results. Furthermore, since the dimensionality of the latent space is much higher than two (10 in (22) and 20 in (23)), those algorithms often rely on external dimension reduction algorithms such as UMAP (24, 25) for visualization functionality, contradicting the premise of a unified probabilistic model.

The competing demands on the model’s latent representation of the data posed by various analysis tasks is hard to balance for a VAE with a shallow latent space. On the one hand, batch effect correction and differential analysis require realistic data generation based on a high-fidelity generative model that preserves detailed information of expression values, which is usually accomplished via increasing the latent dimensionality (26). On the other hand, effective visualization requires a low-dimensional, interpretable latent space, ideally two-dimensional, that captures global biological variation and separates relevant cellular subpopulations. Although a VAE can produce a compact two-dimensional embedding, as reported in (27), this would yield significantly worse synthetic samples than a higher dimensional latent space would. What’s more, VAEs may suffer from posterior collapse (28), particularly when coupled with powerful decoders for better generation capacities. This issue leads to a latent representation that’s indistinguishable from an isotropic Gaussian, rendering interpreting biological heterogeneity such as different cell types almost impossible.

To reconcile these competing objectives, CytoOne adopts a Bayesian hierarchical architecture inspired by Nouveau Variational Autoencoder (NVAE) (20), a powerful deep generative modeling framework that simultaneously supports generation of realistic synthetic data and interpretable low-dimensional representations due to the introduction of multiple levels of latent variables (Fig. 1a). This design enables the model to assign global variation such as cell types to top latent layers for visualization and clustering, while reserving the lower latent layers to model fine-grained variability critical for batch correction and differential analysis. As a result, CytoOne is able to simultaneously learn a two-dimensional representation suitable for exploration, while maintaining sufficient capacity to accurately reconstruct and simulate CyTOF expression profiles.

**Fig. 1.**
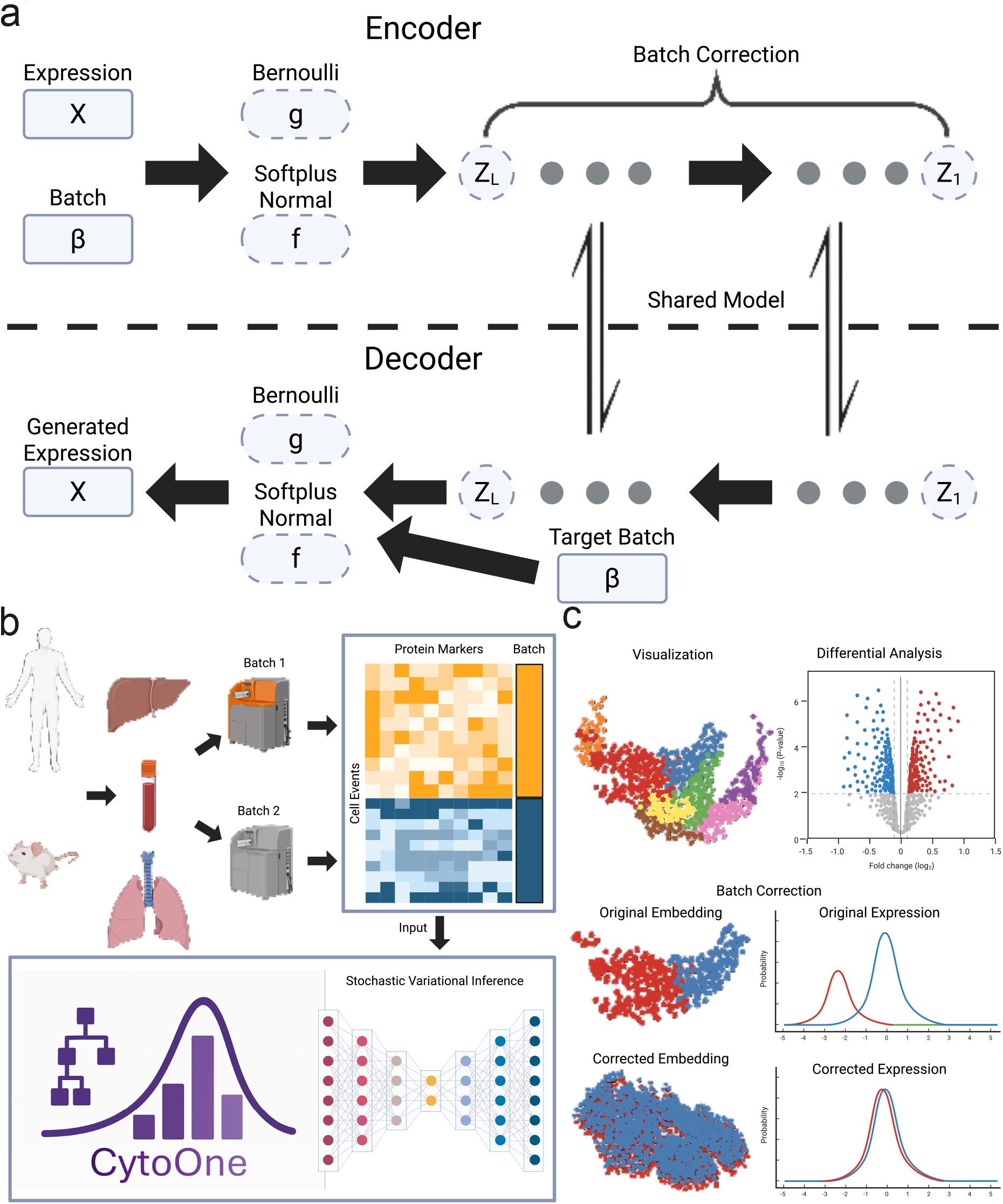
Overview of CytoOne model. (a) The model structure based on NVAE with shared models between the encoder and the decoder. Variables with solid lines indicate observed quantities while variables with dashed lines indicate latent variables. (b) The input and training pipelines of CytoOne. The biological samples can be from different species and various tissues. The inputs are in the forms of expression matrices and their associated batch information. The training process is accomplished via stochastic variational inference. (c) The downstream analyses enabled by CytoOne including direct visualization, differential analysis, and batch correction.

To accurately model the distributional characteristics of CyTOF data explored in (5), we further introduce a novel likelihood function called quasi-zero-inflated softplus-normal distribution (QZIPN). Empirically, CyTOF measurements exhibit zero-inflation or near-zero-inflation where measurements are clustered near zero but not exactly zero. This phenomenon arises from the combination of true biological absence of proteins, contributing to the presence of abundant zeros in the raw output, and the subsequent preprocessing step in the Helios system that adds noise to the original measurements (6). The QZIPN distribution consists of three components: 1. a softplus-transformed Gaussian component that models continuous non-negative expression; 2. a learnable binary gate that models the zero-inflation in the raw output; and 3. a Gaussian convolution kernel to account for added noise. Mathematically, to generate a random variable *x* from a one-dimensional QZIPN(μ, σ^2^, ρ, τ^2^), one can simply carry out: *y* ~N (μ, σ^2^), *f* = log(1 + exp(*y*)), *g* =Bernoulli(ρ), *e* ~N(0, τ^2^), and *x* = *f* ∗ *g* + *e*. By setting τ^2^ = 0, the model will generate zero-inflated expressions.

To further justify our choice of the softplus-normal distribution for modeling CyTOF expression, we performed a direct comparison with the more commonly encountered log-normal distribution (Supp. Fig. 1) based on the gradient of the 75th percentile with respect to distributional parameters. The magnitude of these gradients reflects how rapidly the predicted expression levels change in response to small parameter updates which indicates numerical stability and robustness during training (Supp. Fig. 1a) and generation (Supp. Fig. 1b). The comparison reveals that the softplus-normal distribution offers greater numerical stability across training and generation phases, making it a better fit for CytoOne, where stability is critical for model training and sample generation (See Methods section for details). As the proposed distribution is not of a standard distribution such as the Gaussian distribution or the Binomial distribution, we also validated that a variational inference scheme can be used to approximate the distribution (Supp. Fig. 2a-d). In Supp. Fig. 2e, we also showed that the underlying zero-inflated distribution can be recovered in the presence of added noise (See Methods section for details).

Due to various factors unrelated to biological signals such as sample collection, staining, etc., protein expressions measured from similar systems could exhibit systematic differences (12). Such problems, if not taken care of, would significantly confound the analysis, leading to erroneous conclusions. To address this issue, CytoOne incorporates the batch information and a maximal mean discrepancy (MMD) (29) for penalizing systematic differences in the latent layers between experimental runs. Note that the MMD penalty is only added to latent representations but not to reconstructed protein expressions, which informs the model to separate the underlying biological structure from artificial variations.

The input to CytoOne is an *N* ∗ *M* expression matrix *X* containing *N* single-cell measurements on *M* protein channels. The samples can originate from various tissues of different species. Samples from different batches are simply stacked vertically with an additional column containing β_*n*_ ∈ {1, ···, *B*} indicating the batch information for the *n*th cell event, where *B* is the total number of batches within the data (Fig. 1b). Conforming to the usual CyTOF data analysis workflow (30), we further assume that the protein expression profiles are arcsinh-transformed with a cofactor of 5.

After training, CytoOne outputs two main components. The first component is a batch-effect-free two-dimensional embedding of the CyTOF dataset derived from the top-level latent variables. This is suitable for direct visualization and downstream tasks such as clustering. The second component is the reconstructed/generated protein expressions, sampled from the decoder conditioning on the learned latent space. This output can be used for generating expressions in a target batch which is useful for batch correction and differential expression analysis (Fig. 1c).

### CytoOne well approximates CyTOF data

We first evaluated the extent to which CytoOne can be used as a probabilistic model for approximating the data generating process of CyTOF data by its ability to reconstruct the input protein expressions. To demonstrate the versatility of CytoOne, we used two publicly available datasets: CyAnno (31) which contains raw protein measurements and thus exhibits pronounced zero-inflation, and Levine32 (2, 32, 33) which contains added noise from the aforementioned preprocessing step in the Helios system. After training CytoOne to convergence, we performed ancestral sampling from the generative model to obtain reconstructed protein expression values.

Given the heterogeneous nature of CyTOF data, we began by comparing the observed versus reconstructed expressions within individual cell types. For the sake of demonstration, we focused on Naive CD8+ T cells in CyAnno (Fig. 2a) and CD8 T cells in Levine32 (Fig. 2b) examining a subset of well-characterized marker proteins known to be highly expressed in these populations (34). As negative controls, we also included markers typically expressed in other cell types, such as CD19 (B cells) and CD4 (CD4 T cells).

**Fig. 2.**
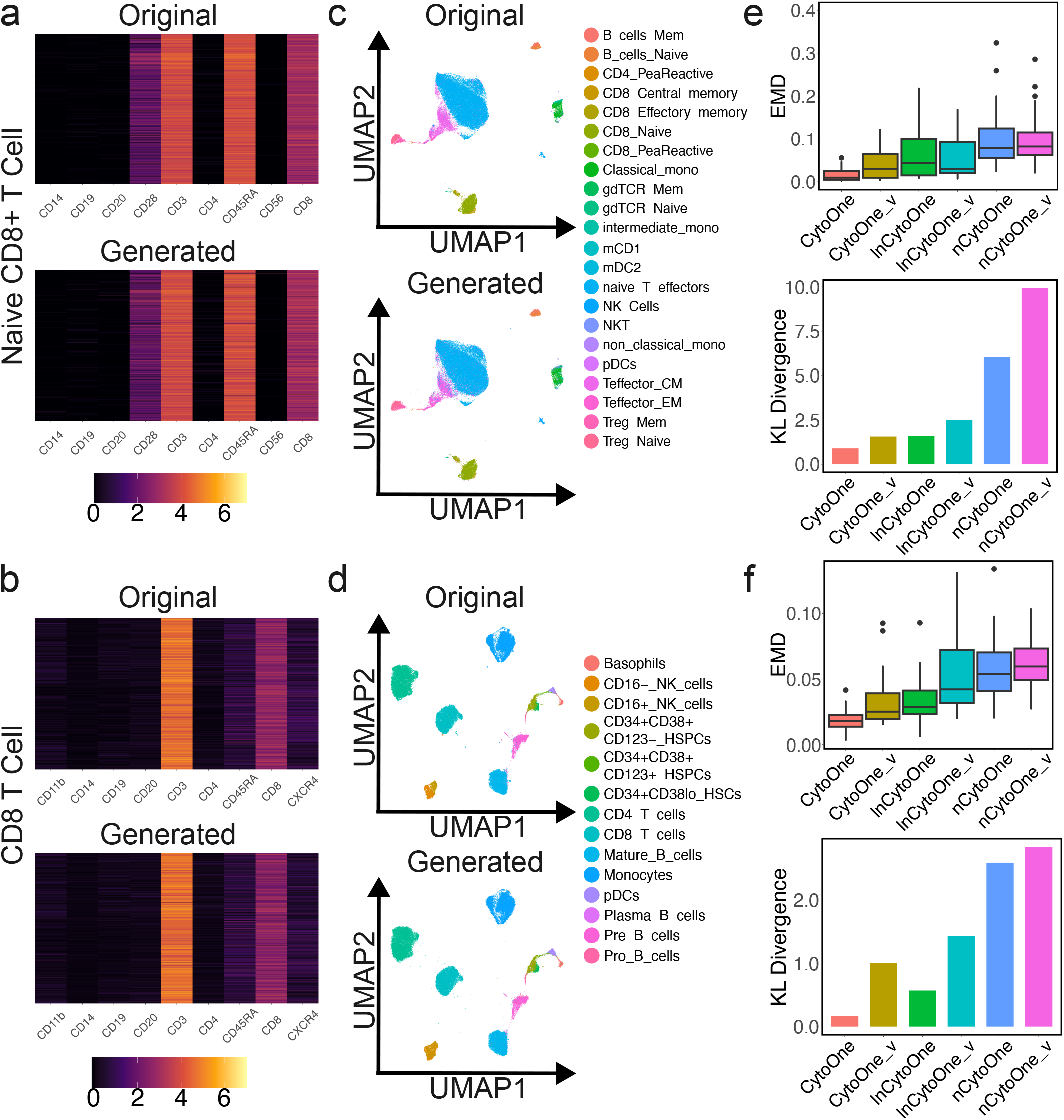
CytoOne well approximates CyTOF data. (a) Heatmaps comparing the expression patterns between original and generated datasets on Naive CD8+ T cells in the CyAnno dataset. (b) Heatmaps comparing the expression patterns between original and generated datasets on CD8 T cells in the Levine32 dataset. (c) UMAP embedding comparing the overall shapes of distributions between the CyAnno dataset and generated dataset. (d) UMAP embedding comparing the overall shapes of distributions between the Levine32 dataset and generated dataset. (e) Quantitative comparisons using EMD and KL divergences among CytoOne and baseline models trained on the CyAnno dataset. (f) Quantitative comparisons using EMD and KL divergences among CytoOne and baseline models trained on the Levine32 dataset.

In both datasets, we observed that the reconstructed expression heatmaps closely mirrored the original data, with minimal distortion. Highly expressed markers such as CD3, CD8, and

CD45RA (for naive T cells) retained their expression profiles in the generated data, while negative control markers were appropriately suppressed—i.e., their reconstructed values were near zero. This suggests that CytoOne effectively captures the marginal distributions of protein expression across biologically relevant subsets of the data. While only two cell types are visualized here for clarity, similar results were observed for other populations and markers.

We then interrogated CytoOne’s ability to approximate the data distribution as a whole by visualizing the dataset using UMAP. As shown in Fig. 2c (CyAnno) and Fig. 2d (Levine32), the global structure of the data is well preserved: the geometric arrangement of cell types, as well as their relative densities and boundaries, remain consistent between observed and reconstructed data. This structural concordance reinforces the validity of CytoOne as a generative model capable of capturing both local expression fidelity and global geometry in CyTOF data.

To complement the qualitative comparisons, we quantitatively assessed how well CytoOne and baseline models approximate the underlying data distribution on both the CyAnno and Levine32 datasets. Specifically, we evaluated marginal distribution fidelity using the Earth Mover’s Distance (EMD) (35) between the observed and generated distributions for each protein marker, and assessed global joint distribution similarity using the Kullback-Leibler (KL) divergence (36).

We benchmarked CytoOne against five alternative generative models:

1. CytoOne_v: a VAE using the same quasi zero-inflated softplus-normal likelihood as CytoOne
2. lnCytoOne: CytoOne with a quasi zero-inflated lognormal likelihood instead of QZIPN
3. lnCytoOne_v: a VAE using the same quasi zero-inflated lognormal likelihood as lnCytoOne
4. nCytoOne: CytoOne with a normal likelihood instead of QZIPN
5. nCytoOne_v: a VAE with a normal likelihood

Across both datasets (Fig. 2ef), CytoOne achieved the best performance on both EMD and KL divergence, demonstrating its ability to recover both fine-grained expression profiles and the overall data structure. CytoOne_v ranked second, showing that even without hierarchical structure, using the right likelihood confers a notable advantage. The performance gap between CytoOne and CytoOne_v, however, highlights the added value of NVAE’s latent hierarchy. nCytoOne with a normal likelihood ranked second to last, reinforcing that the QZIPN distribution is key for modeling CyTOF marginals accurately. nCytoOne_v with a VAE performed worse than its hierarchical counterpart, reflecting both architectural and distributional limitations. lnCytoOne and lnCytoOne_v ranked in the middle, reflecting the advantages brought by quasi zero-inflation likelihood and the drawbacks resulting from the numerical instability of the log-normal component.

Together, these results highlight that the superior performance of CytoOne is largely attributable to two factors: (1) the hierarchical latent architecture from NVAE, which greatly enhanced the capability of CytoOne of generating realistic samples, and (2) the quasi zero-inflated softplus-normal likelihood, which is particularly tailored for modeling the CyTOF measurements. By combining the generation capability of a hierarchical latent representation and a likelihood function tailored for CyTOF data, CytoOne yields a significant performance boost on approximating the structure of CyTOF data.

### CytoOne removes artificial effects

Despite the high-throughput capability of the CyTOF experiment, when the sample size is large, researchers often resort to profiling all samples in separate experiments or batches. As mature as the CyTOF technique, due to differences in sampling procedures, stimulation methods, library preparation protocols, staining techniques, etc., various artifacts that are unrelated to the biological signals of interest could contaminate the obtained protein expressions (11, 12). The resulting artificial variations even in homogeneous samples could potentially jeopardize subsequent downstream analysis.

What’s more, the underlying mechanism through which the distortion inflicts batch effects upon CyTOF samples is rather complicated. In (12) and in (5), they found complex interactions among batches, protein markers, and cell types. To remedy the adverse consequences of batch effects, several methods have been proposed, among which CytofBatchAdjust (13) and CytoNorm (12) enjoy the highest popularity and have been incorporated into data analysis pipelines (16, 17).

CytofBatchAdjust operates by scaling each protein marker to the reference without considering the heterogeneity brought forth by different cell types. On the other hand, CytoNorm alleviates batch effects through a pipeline workflow in which cell clusters are first obtained by FlowSOM (18) by assuming that even in the presence of batch effects, clustering is relatively unaffected. Then, a series of splines are fit for each cluster to normalize samples to the reference sample.

CytoOne entertains a completely different philosophy by jointly modeling batch effects and cell clustering. This is based on the observation that if the data are multi-modal, then the KL-divergence is lower to have a Gaussian distribution for each mode than if the model fits a single Gaussian for all data (37). Therefore, cell clustering is done implicitly rather than requires a separate step. By further informing CytoOne of the batch information and introducing a maximal mean discrepancy (MMD) (29) for penalizing systematic differences in the distributions of the latent representations among batches, we encourage the model to remove the extra modes caused by batch and only retain variations related to biological structures in the latent spaces. With the batch effect removed in latent representations, we then add batch information and train CytoOne to reconstruct the original protein expression. This reflects our understanding that the underlying structure among batches should be similar and batch effect should only manifest at the very last step. To obtain a batch-corrected protein expression matrix with CytoOne, we only need to further inform CytoOne of the index of the reference batch to which we desire to normalize all samples.

To showcase the benefits of joint modeling, we first carried out a simulation study via Cytomulate (5) where a total of 100 pairs of synthetic CyTOF datasets were generated under two scenarios: high variance and low variance. Specifically, for each pair, we first fit Cytomulate using the Levine32 dataset (32). Then, two synthetic datasets representing data from two batches were simulated. While we kept the first batch as is, the batch effects on different cell types in the second batch were simulated via adding an isotropic Gaussian noise with variance σ^2^ and mean being sampled from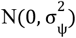.

In the high-variance setting, 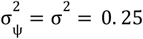 while they were set to 0.01 in the low-variance case. Under each setting, we repeated the process 50 times. After applying all three methods to these datasets, we followed the batch correction benchmark study carried out in (5) and measured the extent to which the batch effects have been attenuated by utilizing the Earth Mover’s Distance (EMD) (35) between two batches. A smaller distance corresponds to a more effective batch normalization.

As expected, when the effects are severe under the high variance setting, the two batches are more dissimilar as compared to the low variance setting (Fig. 3a). Strikingly, under both circumstances, CytoOne achieved the best performance in removing batch effects by a large margin, highlighting the benefits of joint modeling over a pipeline workflow. We further applied three methods to the CytoNorm dataset provided in (12) with three batches, computed EMD after batch correction, and observed a similar result (Fig. 3b).

**Fig. 3.**
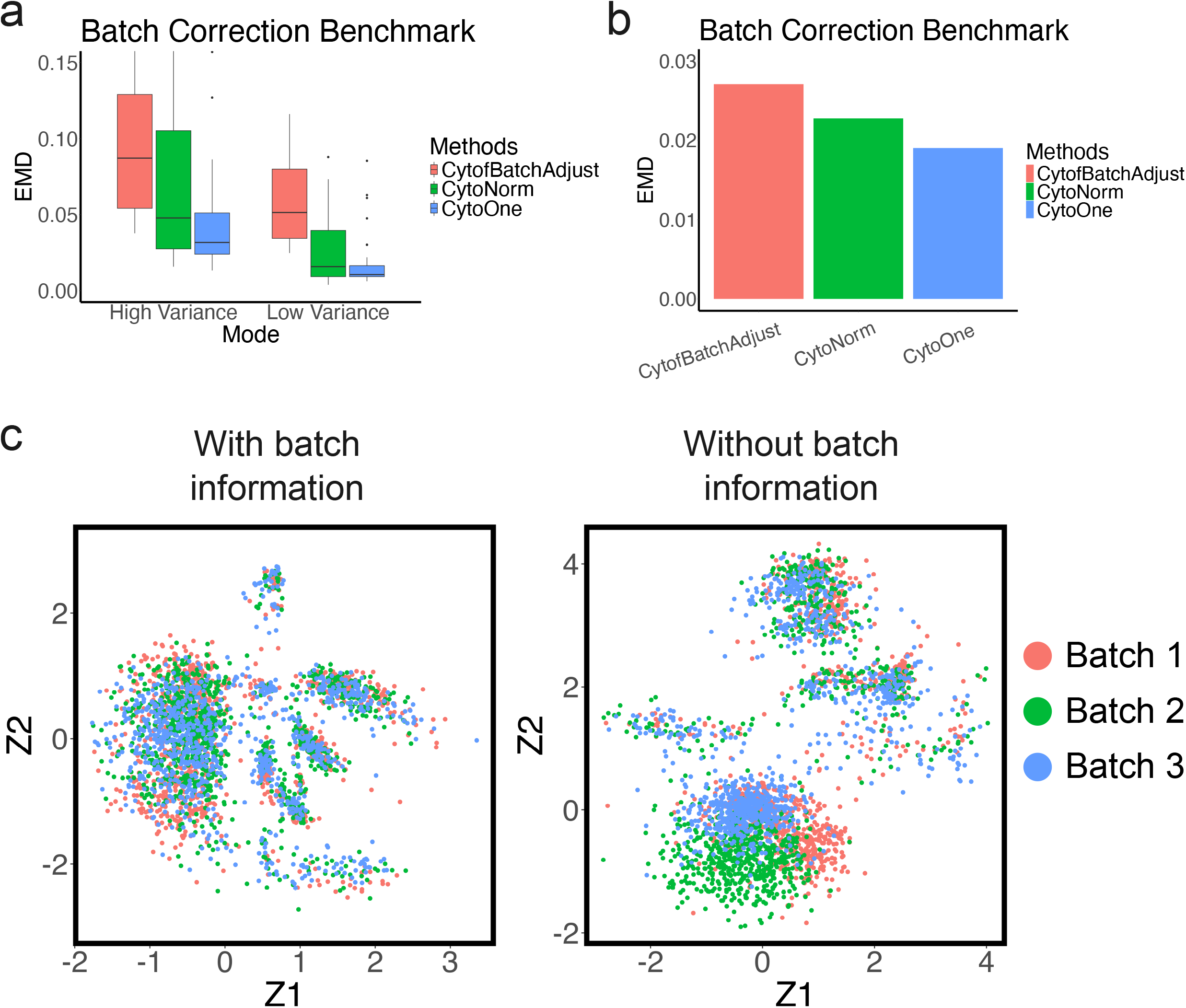
CytoOne removes artificial effects. (a) Batch effect removal results on the simulated datasets generated by Cytomulate. The EMD values are computed by averaging over all protein channels. A lower EMD value indicates a better batch correction. (b) Batch effect removal result on the CytoNorm dataset. The EMD values are computed by averaging over all protein channels. A lower EMD value indicates a better batch correction. (c) An ablation study showcasing that CytoOne effectively utilizes the batch information to remove batch effects in the latent space. The dots are colored according to their batch annotations.

We further conducted an ablation experiment (Fig. 3c) using the CytoNorm dataset to confirm that the batch information has been exploited by CytoOne by training CytoOne without the batch annotation. Notably, when comparing the constructed latent space given the batch annotation with the one rendered without batch annotation, we see that three batches have clear distinctions without the batch annotation (Fig. 3c right) while they are almost indistinguishable from each other given such information (Fig. 3c left). This striking difference confirms CytoOne’s ability to remove extra modes induced by experimental conditions rather than true biological signals.

### CytoOne facilities differential discoveries

To link phenotypes of interest and genotypes under examination, single-cell data analysis often applies differential discovery techniques to compare expressions of cells under two conditions. The analysis is usually carried out cell-type wise based on the measured single-cell resolution expressions. By utilizing distributional information calculated based on those observations such as average, median, and variance, etc., the discrepancy between the measurements among cells of the same type across different conditions are then examined (16). This is often followed by scrutinizing detected differentially expressed protein by field experts via a series of experiments.

The most commonly adopted algorithm in CyTOF differential analysis is diffcyt (15) where the median expression of each protein marker from every sample is computed cell type by cell type, after which adjusted p-values are generated based on the results of a series of linear regressions fit to median expressions. Since the computation is based on summary statistics, in spite of the high-throughput nature of CyTOF data, diffcyt is blazing fast, making it a popular choice in data analysis pipelines.

On the other hand, by constructing a coherent probability distribution over CyTOF data, CytoOne can easily perform differential analysis by sampling from inferred posterior distributions without resorting to summary statistics. This facilitates fully exploiting information in the data and can potentially detect changes on a minute scale.

We first applied both methods to the PBMC dataset (38) with a known differentially expressed protein pS6. For CytoOne, we computed the Bayes factor (39) approximated using posterior samples of the normal component of the QZIPN distribution similar to the procedure described in (22) (See Method section for details).

When we compare the results for different cell types (Fig. 4a), we see that the two techniques yield similar outcomes: both of them found that the protein pS6 is differentially expressed on B cells IgM+ and surface-. By utilizing the posterior samples, CytoOne can further construct posterior distributions for the log-2 fold change of pS6 expressions for each cell type, providing uncertainty quantification beyond the Bayes factor (Fig. 4b).

**Fig. 4.**
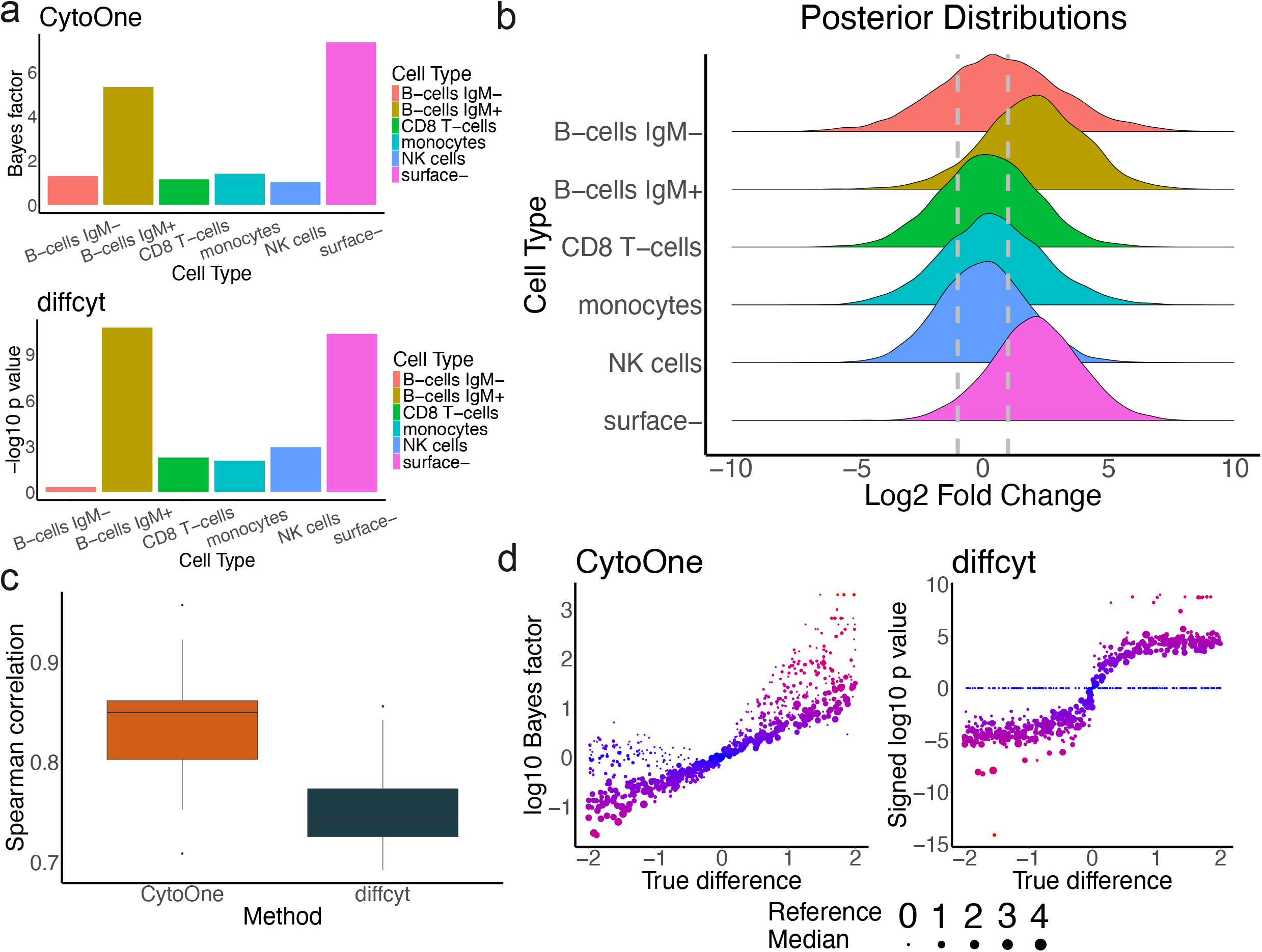
CytoOne facilitates differential discoveries. (a) Differential analysis results on the PBMC datasets across different cell types. (b) Posterior distributions of log-2 fold changes of pS6 expressions for all cell types. Two vertical lines indicate log-2 fold changes of 1 and −1 which are commonly used cutoffs for differential analysis. (c) Differential analysis results on the simulated datasets generated by Cytomulate. The spearman correlations on CytoOne are computed between the true difference and the Bayes factor transformed by the logarithm with a base of 10. The spearman correlations on diffcyt are computed between the true difference and the p-values transformed by the logarithm with a base of 10 and signed by the sign of the true difference. (d) Detailed results on the logarithm-transformed Bayes factor and the signed logarithm-transformed p-values.

Due to the lack of ground truth in the real data, we further conducted a simulation study with 20 sets of synthetic datasets based on the G2 cell type in the brain dataset (40) using Cytomulate (5). Within each set of synthetic datasets, four datasets were generated where the first two representing two replicates under reference condition were retained as is. The condition effect for each protein marker was simulated from a uniform distribution U[−2, 2] and added to the other condition. After applying both methods to the simulated data, we computed Spearman’s correlation (41) between the actual magnitude and the p-values/Bayes factors. The rationale behind this is that the outcomes reported by a differential discovery method should be monotonically correlated with the actual difference.

In Fig. 4c, we see that the Bayes factors returned by CytoOne are overall more correlated with the actual differences. The underlying reason can be illuminated when we examine the result in detail. In Fig. 4d where each dot stands for a protein marker, the colors represent the absolute values of log10 p-values/Bayes factors with red being of higher magnitude and blue being close to 0. The size of a dot shows the median expressions of a protein marker in the reference condition. Since diffcyt solely relies on the median expressions between two conditions, when the data is severely zero-inflated, i.e. when more than 50% of the expressions are 0, no difference can be detected. On the other hand, as CytoOne draws conclusions based on all observations, it is able to detect some difference under such circumstances. This observation corroborates utilizing the entire data distribution as in CytoOne instead of drawing conclusions based on summary statistics.

### CytoOne captures biological variations in its latent space

A critical step in exploring CyTOF data is data visualization achieved by some dimension reduction algorithms such as UMAP (29, 41) and tSNE (17, 42). Based on the results of dimension reduction, cell types can then be assigned, usually via an iterative process of automatic clustering in conjunction with manual gating (21, 22). This is arguably one of the most critical steps in CyTOF data analysis or in general single-cell data analysis as most subsequent modeling and investigation such as cellular trajectory inference and differential discovery are often based on the cell type information obtained at this stage.

Since the original derivation of the classic linear dimension reduction algorithm, Principal Component Analysis (PCA) (43), numerous algorithms have been proposed to address various aspects of shortcomings of PCA. With the prevalence of single-cell data, those methods have also been widely incorporated into data analysis pipelines to capture biological structures.

However, due to the complicated biological nature of single-cell data, to date, there is no one-size-fits-all method that dominates all other tools in all scenarios (13). For example, in (13), they found that when applied to CyTOF data, the dimension reduction tools are highly complementary to each other. What’s more, popular methods such UMAP and tSNE which have been commonly adopted for scRNA-seq data are not necessarily the optimal choice for CyTOF based on different criteria.

In this section, we demonstrate that CytoOne which is tailored for CyTOF data can be a competitive tool to be added to the arsenal of exploratory data analysis of CyTOF. We selected 5 top performers based on the benchmarking results of (13): UMAP (29, 41), tSNE (17, 42), SQuadMDS hybrid (15), SAUCIE (14), and PCA (43). Four datasets with cell type information: brain (44), CyAnno (35), Levine13, and Levine32 (36, 37) are utilized to train CytoOne and the posterior samples of the top-layer latent variable are then directly used as the dimension reduction results.

In Fig. 5a and Supp. Fig. 3, we showed all embedding results and further dyed the points according to the provided cell type information to visually inspect the concordance between the clusters in the latent space and the cell typing results.

**Fig. 5.**
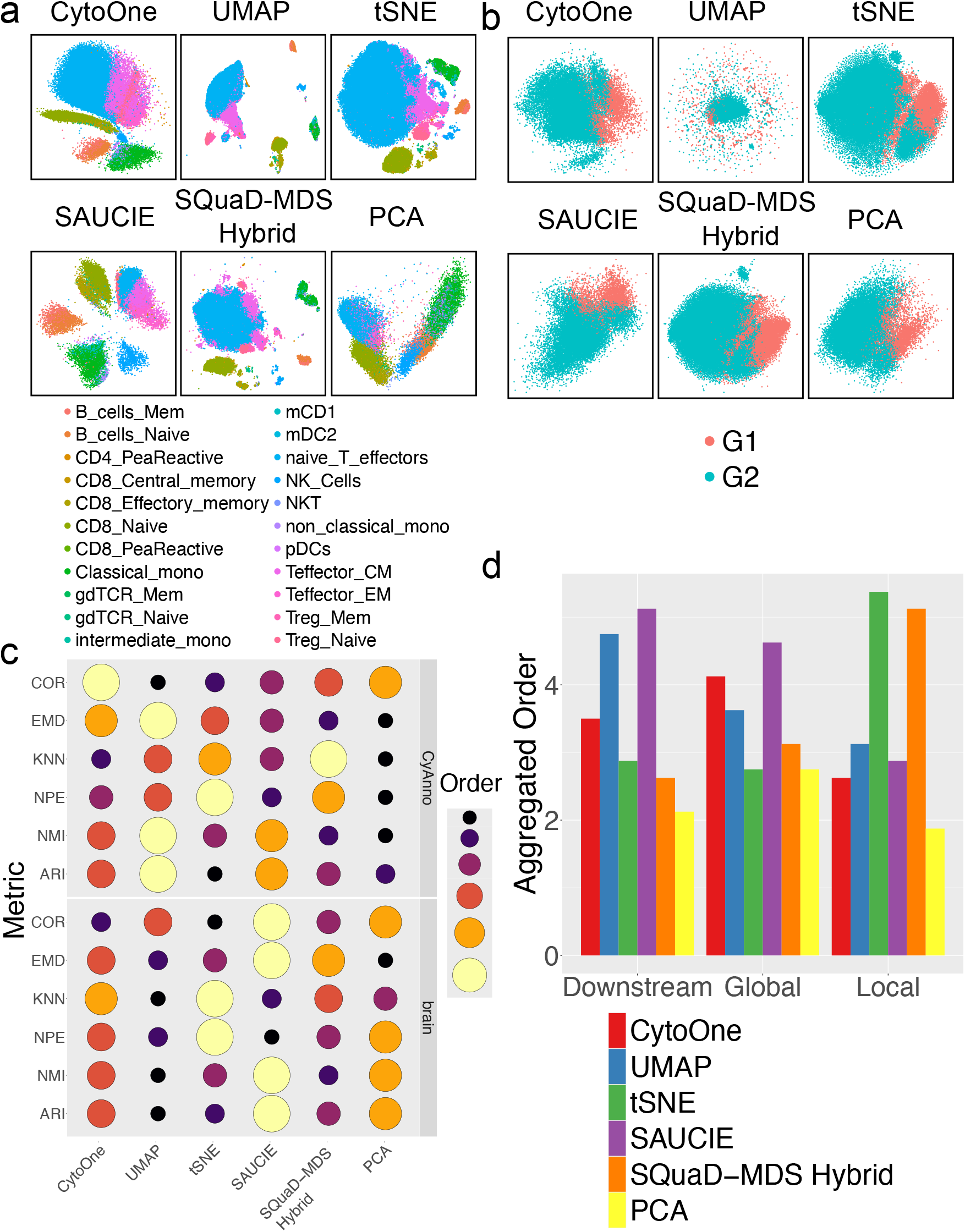
CytoOne captures biological variations in its latent space. (a) Latent embeddings of the CyAnno dataset generated from CytoOne, UMAP, tSNE, SAUCIE, SQuaD-MDS, and PCA. The dots are dyed according to their cell type annotations. (b) Similar to (a) but with latent embeddings of the brain dataset. (c) Quantitative comparisons among the six methods using COR, EMD, KNN, NPE, NMI, and ARI. The size of the dot is proportional to the order of a method on a metric with a higher order indicating a better performance. (d) Aggregated results comparing the six methods. The values are computed by first averaging the orders across all four datasets and then averaging over the categories the metrics belong to.

Across all datasets tested, all non-linear dimension reduction methods seem to be capable of differentiating cell types as cell events of the same type tend to cluster together. One notable exception is UMAP in the brain dataset (Fig. 5b), which produces an odd embedding with cells splattered around the latent space. On the other hand, cell types are less distinguishable based on the results of PCA owing to its linear nature (e.g. Supp. Fig. 3ab). It’s worth noting that during the training stage, all methods are agnostic of the manually gated cell type information. Therefore, the visual clustering in the latent space is formed completely based on the patterns existing in the protein expressions.

To quantitatively investigate the performance of different algorithms, we exploited the tool developed by (13) and computed 6 metrics on the quality of the dimension reduction results: Spearman’s correlation (COR) (45), Earth Mover’s Distance (EMD) (39), K-Nearest neighbors (KNN) (46), Neighborhood proportion error (NPE) (47), Adjusted Rand Index (ARI) (48), and Normalized Mutual Information (NMI) (49). COR and EMD aim to gauge, on a large scale, how well the latent space preserves the original structure in the high-dimensional space. KNN andNPE, on the other hand, assess if the neighboring relationship is retained during the dimension reduction process. Finally, since one of the key functionalities of dimension reduction is to facilitate cell typing, ARI and NMI are adopted to measure the concordance between the actual cell types and the latent space clustering results. In general, COR and EMD are measures of global structure preservation, KNN and NPE focus on local structure preservation, and ARI and NMI examine downstream analysis performance.

We also ordered the six methods based on calculated values of each metric, where the best methods received an order of 6 with the worst being 1. From Fig. 5c and Supp. Fig. 3c, we see that the performance of different methods on various datasets on the six metrics are rather mixed where CytoOne is often ranked in the middle. In Fig. 5d, we further aggregated the results across all four datasets by averaging the orders within each category: downstream analysis, global structure preservation, and local structure preservation. We see that the local structure preservation performance of CytoOne has some weakness. However, this is expected due to the stochastic nature and the regularization by the KL-divergence during the training and inference processes of CytoOne. In other words, instead of aiming at finding a latent representation that can preserve the distance matrix as faithfully as possible, CytoOne, while allowing a meaningful sampling and promoting a structured probabilistic representation, performs denoising and smoothing guided by a Gaussian distribution. This will render the latent space more continuous whilst inevitably slightly distorting the concordance between the representation and the input data.

However, when examining global structure preservation and downstream analysis, we see that CytoOne’s performance is comparable with the top performing dimension reduction methods for CyTOF data reported in (13). Specifically, its global structure preservation surpasses that of UMAP and is almost comparable to that of SAUCIE. Given that interpreting the overall structure of the data such as spatial relationships among clusters and facilitating cell typing are two of the most critical usages of dimension reduction algorithms in CyTOF data analysis, we are confident that CytoOne will be a valuable tool in this endeavor.

## Discussion

The common way to carry out exploratory data analysis for CyTOF data involves a pipeline of various methods with potentially inconsistent or erroneous distributional assumptions, which could result in biased output and often only utilizes partial information contained in the dataset. In the presence of complicated interactions among different facets of data, this sequential fashion of workflow is usually only capable of alleviating the issue moderately. The lack of a unified framework for modeling all features of CyTOF data jointly warrants a computational tool tailored specifically to incorporate unique characteristics of CyTOF data. Once trained, such a model would carry the capacity of performing multiple tasks simultaneously by fully exploiting information in the data.

In this paper, we filled in the gap by proposing CytoOne, a unified probabilistic framework for CyTOF data. It draws on the strength of Bayesian hierarchical models (39) and the recent advancement in deep generative frameworks (19, 20) to depict the underlying data generating mechanism. CytoOne takes as input the arcsinh-transformed protein expressions along with annotations on batches, optimizes the variational lower bound via efficient stochastic gradient descent (21), and produces a well-defined probability distribution over the entire input data. Furthermore, its carefully designed hierarchical structure allows researchers to extract information relevant to their hypothesis with ease.

By thoroughly interrogating every aspect of CytoOne and benchmarking its performance in batch correction, differential discovery, and data visualization, we demonstrated that not only can CytoOne well approximate the true data distribution, but also compares favorably to commonly used data analysis pipelines at the tasks they are designed for, highlighting the benefits of joint modeling.

Powerful as CytoOne is, we can see several future directions along which the model can be improved upon. Since the architecture of the deep neural networks of CytoOne is built based on heuristic choices of layer normalization (42) and GELU (43), as well as the widths of various hidden layers, a large scale model selection and incorporating more advanced network structures could potentially further boost its ability for information extraction. Furthermore, since the latent space is designed for visualization purposes, its top level latent dimension is always set to 2. However, as an integral part of the probabilistic model, the choice of the dimensionality of the latent space intuitively would affect the performance of other aspects of the model such as batch normalization and differential discovery. If visualization is not sought after, then perhaps a quick model selection procedure based on BIC (44) can be incorporated into the model fitting process.

Another aspect of the model that we have yet to explore is its potential to generate synthetic CyTOF datasets. However, due to the well-known prior-hole problem prevalent in VAE (45), direct ancestral sampling starting from Gaussian distribution would surely yield unrealistic samples. As a potential remedy, we conjecture that training a diffusion model (46–49) on the latent variables while “freezing” the rest of the model would close the gap between the isotropic Gaussian and the true distribution over the latent representations. With CytoOne’s current capacity and its potential, we are confident that our probabilistic framework will be of great interests in the research community and inspire further exploration in these exciting directions.

## Methods

### The CytoOne model

We attempt to model the arcsinh-transformed data with a cofactor of 5. Suppose we have *b* = 1, ···, *B* batches, *n* = 1, ···, *N* cell events, *m* = 1, ···, *M* protein markers and *l* = 1, ···, *L* hidden layers in NVAE. We denote by *X* the observed data where the *n*th row *x*_*n*_ ∈ **ℛ**^*M*^ contains the protein measurements on the *n*th cell event. Let the batch of the *n*th cell event be denoted by β_*n*_ ∈ {1, ···, *B*} and η(.) = log(1 + exp(.)) be the softplus function. The data generating mechanism is described as follows:

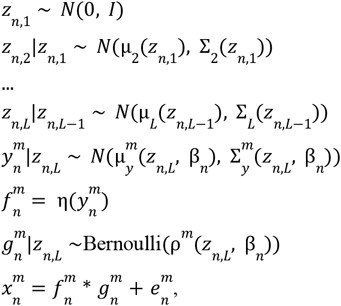

where 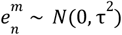 if the data is not zero-inflated and 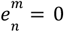 otherwise. μ_2_, ···, μ_*L*_, μ_*y*_, ∑_2_, ···, ∑_*L*_, ∑_*y*_, ρ are all neural networks such that the outputs of ∑_2_, ···, ∑_*L*_, ∑_*y*_ are constrained to be diagonal covariance matrices while each component of the output of ρ is restricted within (0, 1).

For inference, finding a variational distribution for each latent variable is not necessary. Given *z*_*n,L*_, we can integrate out the latent variables 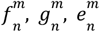 to obtain the marginal distribution 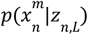. For the sake of clarity, in the following derivation, we abuse the notations a bit and omit the sub- and superscript for brevity. To get started, suppose that given a particular value of *z, y* ~ *N*(μ, σ^2^) and *g* ~Bernoulli(ρ). Then via the change of variable formula, we have 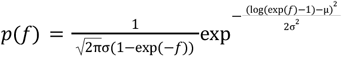, where *f* > 0. Therefore, the distribution of *h* = *f* ∗ *g* is *p*(*h*) = 1 − ρ + ρ*p*(*f*). Finally, when the data is not zero-inflated, we have, 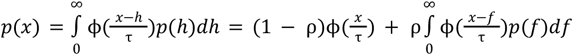. Since it is non-trivial to evaluate this integral analytically, we use Gauss-Hermite quadrature (50) to compute its value. Given the marginal distribution, we can decompose the variational distribution into 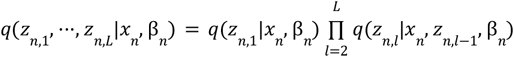 where each component is chosen to be Gaussian with diagonal covariance matrix.

The evidence lower bound (ELBO) can thus be written as follows:

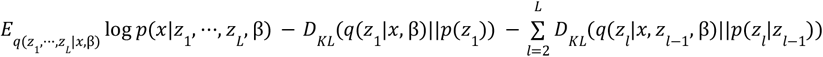

where *D*_*KL*_ stands for the Kullback-Leibler divergence.

To remove batch effect, we further modify the ELBO:

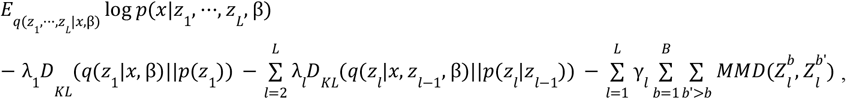

where MMD stands for maximum mean discrepancy and 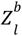 is the latent variables at the *l*th hidden layer resulting from the *b*th batch.

Throughout the paper, we use the Adam optimizer (21) with a learning rate of 1e-3. For the weights on the KL divergences, we set λ_1_ = 0. 01, with the rest being proportional to the latent dimensions. For the weights on the MMDs, we used γ_1_ = 2, with the rest being inversely proportional to the latent dimensions. We used four latent layers with dimensions being 20, 10, 5, and 2. We optimize the objective via stochastic variational inference until convergence, usually around 50-100 epochs.

### Comparison between Softplus-Normal and Log-Normal

In Supp. Fig. 1a, we compared the gradients of the 75% quantile with respect to the location parameter and the scale parameter, respectively. We considered two scenarios: 1. Parameter initialization phase, which is intended to mimic the early phase of model training; 2. Generation phase, which is intended to emulate the model’s behavior during posterior sampling or reconstruction.

In the first scenario, we initialized both distributions (softplus-normal and log-normal) with identical location and scale parameters. We then computed the gradient of the 75th percentile with respect to the location parameter and the scale parameter, respectively. It can be seen from Supp. Fig. 1a, the log-normal exhibited much sharper gradients than the softplus-normal, indicating a high degree of sensitivity to parameter perturbations. Such steep gradients can cause instability in optimization, especially with stochastic gradient descent. In contrast, the softplus-normal demonstrated more moderate and stable gradients, which are more conducive to robust training dynamics.

In the second scenario, we suppose the model has already been trained to fit a target distribution with a fixed mean and variance. By computing the corresponding location and scale parameters of both distributions to match the mean and variance of the target distribution, we again computed the gradients of the 75th percentile with respect to location and scale. We can see from Supp. Fig. 1b, the log-normal remains highly sensitive to small perturbations in the parameters which occur frequently during posterior sampling where the parameters are sampled from another distribution. On the other hand, the softplus-normal maintained consistent gradient behavior.

This gradient-based comparison reveals that the softplus-normal distribution offers greater numerical stability and smoother gradient landscapes across training and generation phases.

### Approximation of QZIPN distributions

To validate that a stochastic variational scheme can be used to approximate a QZIPN distribution, we simulated four sets of QZIPN(μ, σ^2^, ρ, τ^2^) distributions by varying one parameter of the distribution while keeping the others invariant. Specifically, let the baseline configuration be μ = 5, σ = 1, ρ = 0. 5, τ = 0. In the first set (Supp. Fig. 2a), we varied μ from 0.5 to 5 with a 0.5 increment. In the second set (Supp. Fig. 2b), we changed σ from 0.1 to 1 with an increment of 0.1. We increased ρ from 0 to 0.9 with an increment of 0.1 in the third set (Supp. Fig. 2c). Finally, we induced a Gaussian noise with scale τ changing from 0 to 0.9 in the fourth set (Supp. Fig. 2d). In Supp. Fig. 2a-d, the top panels show the original densities while the bottom ones are the density estimates from the posterior samples. It should be evident that the stochastic variational inference scheme we proposed can adequately approximate the QZIPN distribution. In Supp. Fig. 2e, we reused the configuration in Supp. Fig. 2d and extracted the corresponding zero-inflated distributions without noise corruption. By comparing the top panel with the bottom panel, we see that the posterior samples can effectively recover the noise-free distributions.

### Batch effect correction with CytoOne

To perform batch effect correction with CytoOne for the *n*th cell event *x*_*n*_, in addition to its original batch annotation β_*n*_, we need to inform CytoOne of the target batch α_*n*_ to which we desire to normalize the sample. Batch correction for *x*_*n*_ is then carried out according to the following procedure:

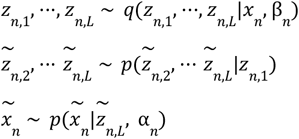

The intuition behind this procedure is that once the encoder removes the batch effect the latent space, the decoder can then map all the latent variables to the target batch.

### Differential analysis with CytoOne

Since CytoOne is constructed via a Bayesian hierarchical model, we computed the Bayes factor (39) approximated using posterior samples of the normal component of the QZIPN distribution *y* similar to the procedure described in (22). Specifically, given a protein marker *m* and the posterior distributions of *y*^*m*,1^ and *y*^*m*,2^ and their corresponding observed protein expressions *x*^1^and *x*^2^ under condition 1 and 2, respectively, we can formulate two hypotheses:

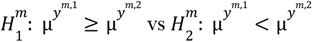

Then, the posterior of 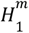 can be approximated via

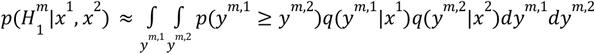

The Bayes factor can then be computed as

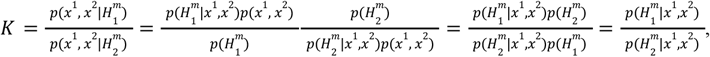

where we assumed that the two hypotheses have equal prior probability: 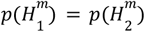. By further sampling expressions from one cell type under two conditions and averaging the obtained Bayes factors, we can obtain an estimate of whether or not the protein marker *m* tends to be more expressed under condition 1 than does condition 2.

### Software implementation details

All analyses, plotting, and benchmarks were performed using the R (4.4.1) and Python (3.10.16) programming environments in MacOS 15.5. CytoOne is implemented and supported in Python (v3.9 or later), and the CLI is available on all platforms and systems where a compatible python interpreter is present with the proper package installations. The detailed documentation and tutorials are freely available and hosted on GitHub (https://github.com/Yuqiu-Yang/CytoOne).

All default settings were used for CytofBatchAdjust, CytoNorm, and diffcyt. For diffcyt, we supplemented the algorithm with cell type information instead of using the clustering algorithm that comes with diffcyt.

### Statistical analyses

For all order-based metrics, higher orders represent better performance. In cases where ties occurred, we used the average order of tied values. For all boxplots appearing in this study, box boundaries represent interquartile ranges, whiskers extend to the most extreme data point which is no more than 1.5 times the interquartile range, and the line in the middle of the box represents the median.

## Declarations

### Ethics approval and consent to participate

Not applicable

### Consent for publication

Not applicable

### Availability of data and materials

The Levine32, Levine13 (32), and PBMC datasets (38) can be accessed from the HDCytoData package (33) in R (https://github.com/lmweber/HDCytoData). The CyAnno (FlowRepository ID: FR-FCM-Z2V9) can be accessed from its publication (31). The brain dataset (40) can be downloaded from https://flowrepository.org/experiments/1734/download_ziped_files. The CytoNorm dataset (12) can be accessed from their GitHub repo: https://github.com/saeyslab/CytoNorm.

The CytoOne algorithm is available at https://github.com/Yuqiu-Yang/CytoOne. All the datasets as well as the source code used to generate the findings are available in Zenodo (https://zenodo.org/records/17795487) and are released under the BSD-3-Clause license.

### Authors’ contributions

Y.Y. performed all analyses. Y.Y. and K.W. created the *CytoOne* Package. X.W. conceived the study. X.W. designed and supervised the whole study. All authors wrote the manuscript.

## Acknowledgements

Not applicable

## Supplementary information

### Additional file 1

**Supp. Table 1:** A table that provides the dataset name, species, the anatomic site, and its source for accession.

### Additional file 2

**Supp. Fig. 1.**
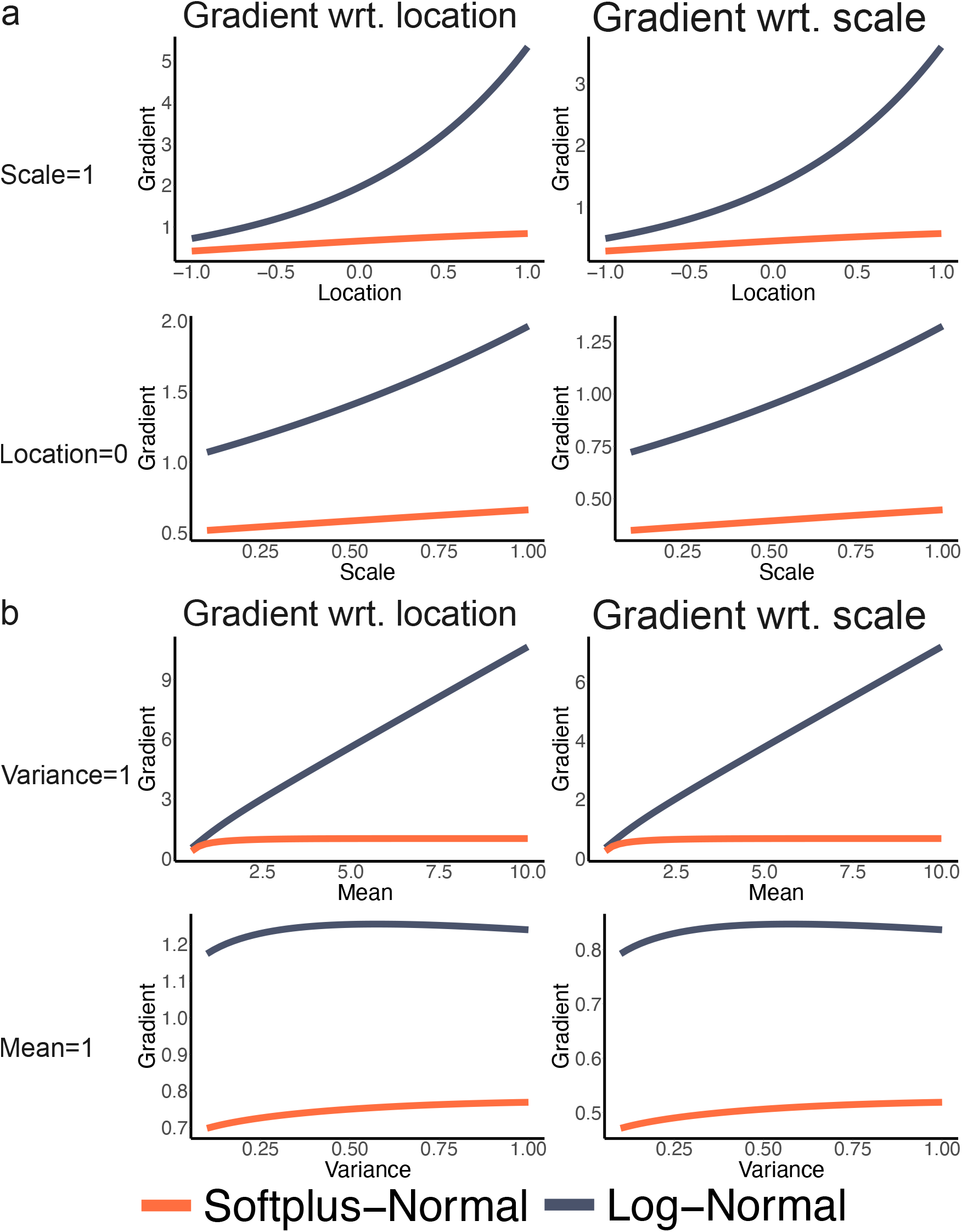
Comparison between softplus-normal and log-normal. (a) Comparison of the gradient of the 75% quantile with respect to the location and the scale parameters between softplus-normal and log-normal by mimicking the parameter initialization phase. The top two panels are computed by varying the location parameter while keeping the scale being fixed at 1. The bottom two panels are computed by varying the scale parameter while keeping the location being fixed at 0. (b) Comparison of the gradient of the 75% quantile with respect to the location and the scale parameters between softplus-normal and log-normal by mimicking the posterior sampling phase. The top two panels are computed by varying the mean of the target distribution while keeping the variance being fixed at 1. The bottom two panels are computed by varying the variance of the target distribution while keeping the mean being fixed at 1.

**Supp. Fig. 2.**
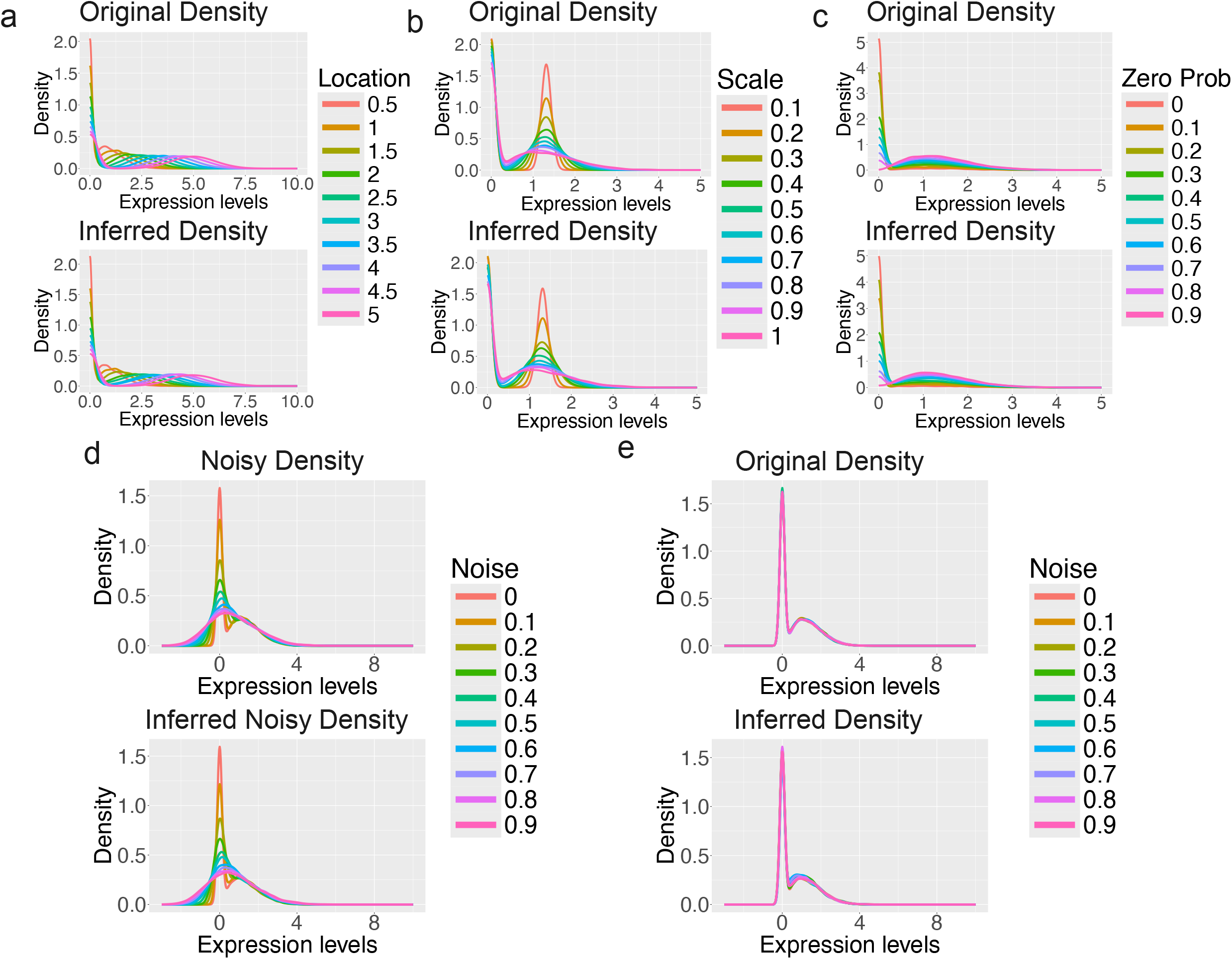
Stochastic variational inference on simulated QZIPN distributions. (a) Density plots comparing the original distributions and the density estimates of the posterior samples. The original distributions are generated by varying the location parameter in the QZIPN distribution. (b) Density plots comparing the original distributions and the density estimates of the posterior samples. The original distributions are generated by varying the scale parameter in the QZIPN distribution. (c) Density plots comparing the original distributions and the density estimates of the posterior samples. The original distributions are generated by varying the zero-inflation gate parameter in the QZIPN distribution. (d) Density plots comparing the original distributions and the density estimates of the posterior samples. The original distributions are generated by varying the scale parameter of the added noise in the QZIPN distribution. (e) Density plots comparing the underlying zero-inflated distributions from the noise-corrupted distributions in (d) and the density estimates of the posterior samples.

**Supp. Fig. 3.**
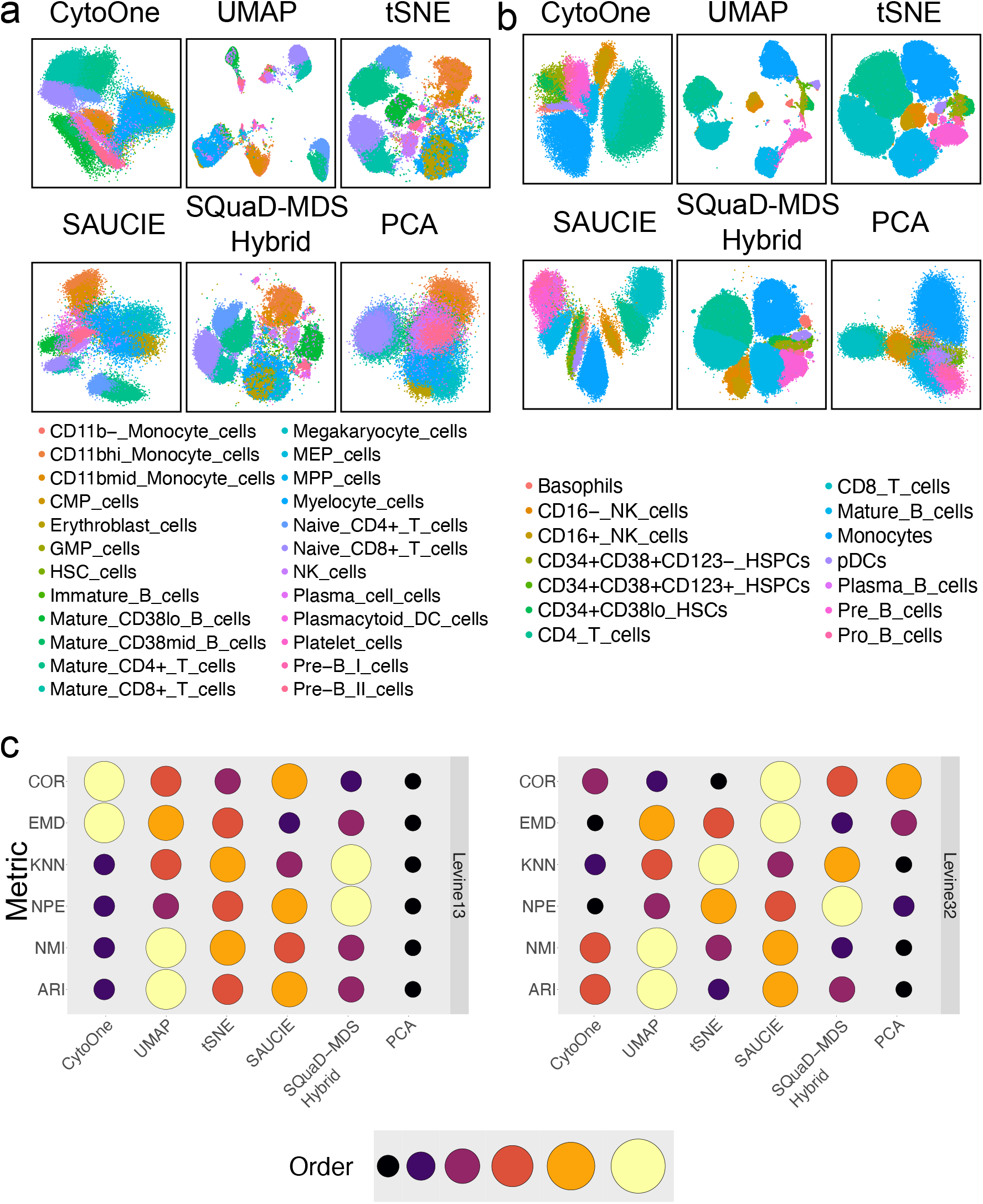
Additional DR comparisons among the six methods. (a) Embeddings generated based on the Levine13 dataset. (b) Embeddings generated based on the Levine32 dataset. All dots are dyed according to their corresponding cell type annotations. (c) Quantitative comparisons among the six methods using COR, EMD, KNN, NPE, NMI, and ARI. The size of the dot is proportional to the order of a method on a metric with a higher order indicating a better performance.

