## Supplementary material for "Unified Probabilistic Analysis of CyTOF: A Deep Generative Approach using CytoOne": Supp. Table 1

| Dataset | Species | Anatomic Site | Source | Number of Markers | Number of Cells Events | Number of Cell Types |
|---|---|---|---|---|---|---|
| Levine32 | Human | Bone Marrow | HDCytoData | 32 | 72,463 | 32 |
| Levine13 | Human | Bone Marrow | HDCytoData | 13 | 167,044 | 24 |
| brain | Human | Brain | https://flowrepository.org/experiments/1734/download_ziped_files | 35 | 49,585 | 2 |
| CyAnno | Human | Peripheral Blood | https://flowrepository.org/id/FR-FCM-Z2V9 | 39 | 123,033 | 39 |
| PBMA | Human | Peripheral Blood | HDCytoData | 33 | 172,791 | 6 |
| CytoNorm | Human | Blood | https://github.com/saeyslab/CytoNorm | 37 | 3,000 | |
